# Phylogeographic and morphological analysis of *Botrylloides niger* Herdman, 1886 from the northeastern Mediterranean Sea

**DOI:** 10.1101/2022.11.30.518487

**Authors:** Berivan Temiz, Esra Öztürk, Simon Blanchoud, Arzu Karahan

## Abstract

*Botrylloides niger* is an invasive marine filter-feeding invertebrate that is believed to originate from the West Atlantic region. This species of colonial tunicate has been observed on several locations along the coasts of Israel and around the Suez Canal but it has not yet been reported on the coasts of the northeastern Mediterranean Sea (NEMS), suggesting an ongoing Lessepsian migration. However, the extent of this invasion might be concealed by reports of other potentially misidentified species of *Botrylloides*, given that the strong morphological similarities within this genus renders taxonomical identification particularly challenging. In this study, we performed a phylogeographic and morphological analysis of *B. niger* in the NEMS. We collected 241 samples from 8 sampling stations covering 824 km of coastlines of NEMS. We reported 14 different morphotypes, of which the orange-brown, orange and brown-striped morphs were the most abundant. Using the mitochondrial cytochrome C oxidase I (COI), one of the four most commonly used DNA barcoding marker, we identified 4 haplotypes with the Konacık (H4) and the Mezitli (H3) ones being the most diverged. The COI haplotypes clustered with the reference *B. niger* sequences from GenBank and separated from sister *Botrylloides* species with high confidence. We confirmed our identification using the three additional barcoding markers (Histone 3, 18S rRNA and 28S rRNA), which all matched with over 99% similarity to the reference sequences. In addition, we monitored the Kızkalesi station for a year and applied temporal analysis to the colonies collected. The colonies regressed during winter while resettled and expanded during summer. We performed gene flow analysis on our spatial data that identified a possible population subdivision at the sampling site of Side, which might be caused by a local freshwater input. Overall, we here present the first report on the presence of *Botrylloides niger* in the NEMS, we show that this species is commonly present throughout this region and with a particularly high morphological as well as genomic diversity.

## 1. Introduction

Phylogeography is a relatively new field that investigates genealogical lineages’ geographical distribution (Avise, 2009). Populations divided by long distances or geographical barriers may display a higher level of genetic variation representing the accumulation of mutations acquired over long periods of isolation (Bowen et al., 2016). Geography is thus intrinsically connected with evolutionary relations, and this connection can be detected via molecular markers that estimate genetic variations to understand the linkage between congeneric species (Avise, 2009). Consequently, spatial analyses of genetic diversity can provide important insights into the evolution of entire populations over potentially large spatial and temporal scales.

While such analyses are reasonably accessible in terrestrial environments, in aquatic systems the population dynamics are more turbulent and harder to estimate. Considering the development in maritime traffic and aquaculture, the establishment of exogenous marine organisms in new environments is inevitable (Seebens et al., 2016). In particular, the global spreading of invasive species is a major threat to endogenous ecosystems and their prevention is of major biological and economical interest. Thus, phylogeographic studies on marine environments are essential tools to estimate gene flow, genetic diversity, isolation patterns, and hydrographic barriers to elucidate the dispersal of populations.

DNA barcoding is a prominent molecular tool used to identify species and catalogue biodiversity. The main marker used for barcoding is a 500-600 bp fragment of the mitochondrial cytochrome C oxidase subunit I (COI) gene, which sequences are compared to global reference databases such as GenBank or Barcoding of Life Database (BOLD) (Ratnasingham & Hebert, 2007; Benson et al., 2012). Additional barcoding markers including sequences of the chromatin component Histone 3 (H3), of the ribosomal small subunit 18S rRNA, and of the ribosomal large subunit 28S rRNA have been used to resolve population dynamics at the sub-species level.

In the Northeastern Mediterranean Sea (NEMS), DNA barcoding studies have increased in recent years to monitor possible Lessepsian migrations from the Suez Canal for controlling the resulting invasions (Karahan et al., 2017). Such researches have proven to be decisive for the implementation of marine conservation policies by providing the required scientific knowledge on the ecosystem’s dynamics (Eryilmaz & Dalyan, 2006; Smith et al., 2008; Keskin & Atar, 2013; Oter et al., 2013; Azzurro et al., 2015; Bariche et al., 2015; Seyhan & Turan, 2016; Tuney, 2016; Ciftci et al., 2017; Karahan et al., 2017; Ozbek et al., 2017; Reem et al., 2018; Golestani et al., 2019).

*Botrylloides* is a genus of sessile marine invertebrate filter-feeders that belong to the Tunicata subphylum and that are present globally (Shenkar & Swalla, 2011). These colonial chordates live on hard substrata by establishing flat and gelatinous colonies after the motile larva attaches and undergoes metamorphosis (Berrill, 1947). *Botrylloides* can also undergo asexual reproduction, which is called blastogenesis, whereby new zooids bud from the peribranchial wall of the parental zooids (Sabbadin, 1958). The blastogenic cycle culminates with the death and absorption of all parental zooids by the colony and the emergence of a new generation of zooids in a process known as the takeover (Manni et al., 2019). Natural chimerism following the fusion of two closely related kins, hibernation and even whole-body regeneration have commonly but not exhaustively been reported in *Botrylloides* (Paz & Rinkevich, 2002; Rinkevich, 2005; Sheets et al., 2016; Zondag et al., 2016; Blanchoud et al., 2017; Blanchoud et al., 2018a; Karahan et al., 2022). The *Botrylloides* genus is composed of 21 reported species, all of which are morphologically very close with zooids aligned in a ladder-like arrangement. Anatomical features that differentiate species can be very difficult to assess, such as the number of stigmata rows on the branchial basket, the presence of a pyloric caecum or the type of larval incubation. Consequently, taxonomical assignments based on a sample’s anatomy only can be insufficient for species-level categorization and misclassifications in the proper identification of *Botrylloides* species have been frequently reported, contaminating even reference databases (Viard et al., 2019).

Several species of *Botrylloides* have been identified as invasive and were reported in the Mediterranean Sea (Spallanzani, 1784; Olivi, 1792; Savigny, 1816; Pinar, 1974; Rinkevich et al., 1993; Brunetti, 2009; Brunetti & Mastrototaro, 2012; Cinar, 2014; Halim & Messeih, 2016; Reem et al., 2018; Viard et al., 2019). *Botrylloides niger* (Herdman, 1886) is classified within the Ascidiacea (Class) and has been synonymized with reports of other colonial ascidians including *Metrocarpa nigrum* (Herdman, 1886), *Botryllus niger* (Herdman, 1886), *Botryllus nigrum* (Herdman, 1886), *Botrylloides chazaliei* (Sluiter, 1898) and *Botrylloides nigrum* (Herdman, 1886). *B. niger* is predicted to be native from the West Atlantic due to its frequent presence there, although it was first identified on the coasts of Bermuda, an island located in the temperate region of the North-Atlantic ocean (Sheets et al., 2016). Peres (1958) documented the presence of *B. niger* on the Mediterranean coasts of Israel over fifty years ago already, and Halim and Messeih (2016) recently reported its presence in the Suez Canal.

The only record of *Botrylloides* in the NEMS is that of *Botrylloides leachii*, documented in Mersin harbor in Türkiye (Pinar, 1974). However, this report is solely based on morphology, and with a number of missing key anatomical insights, which suggests that the resulting taxonomical assignment at the species level could be debatable. For instance, Brunetti (2009) and Reem et al. (2018) disagree with Sheets et al. (2016) on the identification of *Botrylloides* colonies on the coasts of Israel, with the former ones assigning them as *B. leachii* but the latter ones as *B. niger*.

In this study, we investigated the genetic and morphological diversity of *B. niger* colonies from the NEMS by combining COI barcoding with H3, 18S rRNA and 28S rRNA to support our identification, all sequences clustering phylogenetically with the available references for *B. niger*. Through sampling the 824 km coastal range of the NEMS, we described 14 morphotypes and 4 haplotypes, with their life cycle including blastogenesis and hibernation. In addition, we conducted a time-series sampling for a year to investigate the effect of seasonal changes on the population dynamics and show that the colonies are not present during winter but expand during summer. Using gene flow analysis on our spatial data, we measured that the populations were under negative selection and suggest that freshwater input can be a factor for this population division. Overall, we here provide the first large-scale high-resolution documentation of the globally invasive species *Botrylloides niger* in the NEMS.

## 2. Materials and methods

### 2.1. Colonies sampling

Spatial samples were collected from the coastal areas of Antalya (Kemer-Side--Alanya-Tersane), Mersin (Tisan-Kızkalesi-Mezitli), and Hatay (Konacık) in September and October 2018) in water depth between 30-50 cm as previously described for *Botrylloides anceps* (Figure 1A) (Karahan et al., 2022). Samples were collected from the submerged stones using a single-edged razor blade. The colonies to be monitored were taken onto a microscope slide and attached with a sewing thread while the samples for barcoding were put in 1.5 mL tubes filed 70 % (v/v) ethanol. The sampling area’s date, coordinates, salinity, and pH were documented (Table S1). In total, 278 samples were collected, including additional spatial samples (Table S2) and additional samples for the Kızkalesi time-series collected between November 2017 and October 2018 (Table S3).

**Figure 1.**
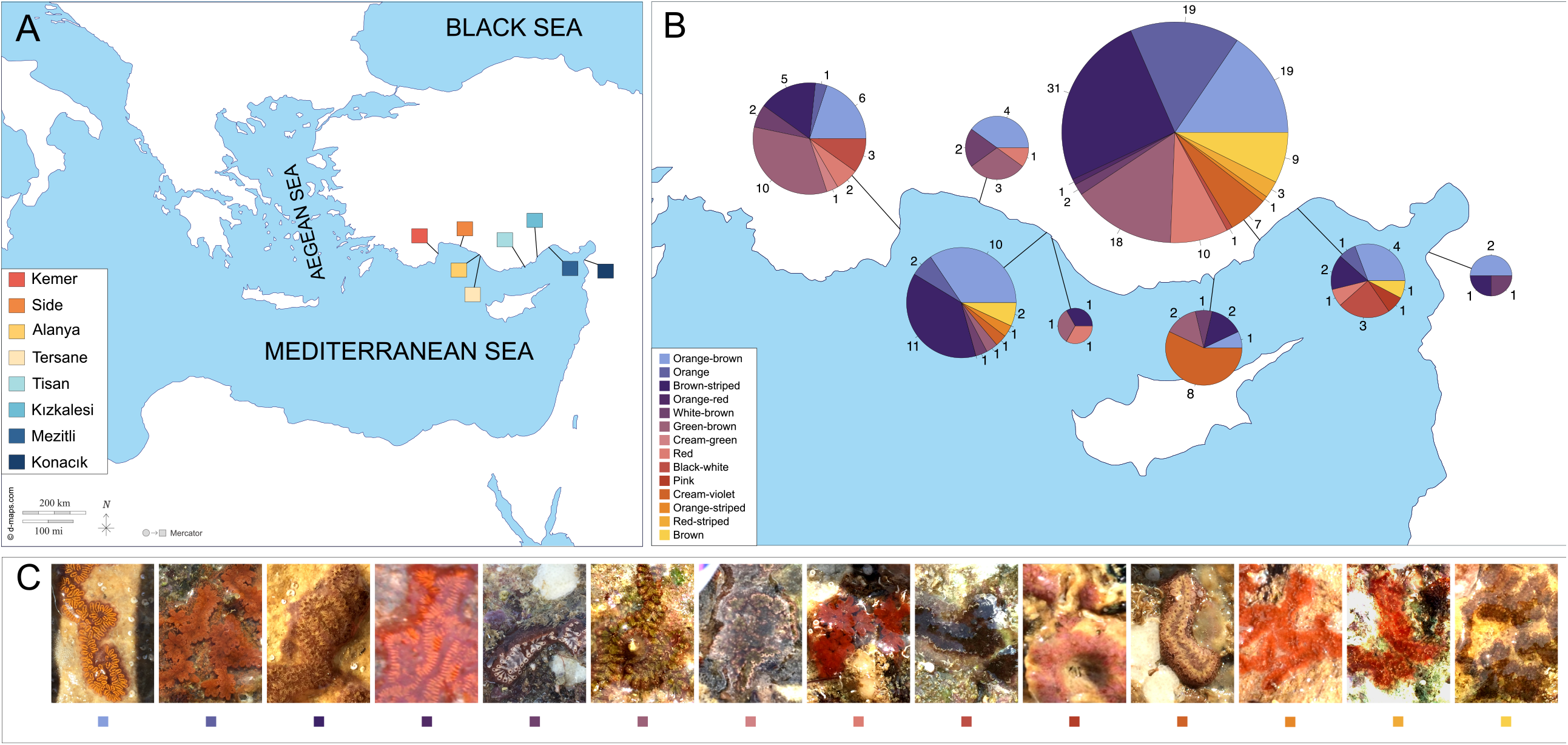
Morphotypes of *Botrylloides niger* from the sampling stations of the NEMS. **A)** NEMS sampling sites, from West to East, cover the Antalya region (Kemer, Side, Alanya, Tersane), the Mersin region (Tisan, Kızkalesi, Mezitli) and the Hatay region (Konacık). **B)** Regional diversity in the sampled morphotypes, depicted with pie charts proportional to the sample sizes. **C)** *In-situ* images of the corresponding morphotypes.

### 2.2. Colonies Characterization

All tunicate specimens were characterized via morphological examination regarding their zooid distribution, colour, and habitat preferences. Living specimens that were taken to the laboratory were photographed under a stereo and a light microscope (Olympus SZX16 - UC30 camera; Olympus CX43-ToupTek camera). Living colonies were kept at 20 °C in the IMS-METU aquaculture room. The temperature and light of the room were stable, and the salinity of the water was constant at 40 ppt. The colonies were fed regularly, and the water was replaced every two days.

Hibernation of the animals was characterized by visual inspection based on the regression of all the zooids, hence resulting in a dense vascular system. Blastogenic cycles were monitored from the ventral side of the zooids and the budding through the atrial epithelium was monitored by visual inspection.

### 2.3. Molecular analyses

DNA extractions were completed based on the protocol of (Karahan et al., 2022). The amplification reactions were executed based on Reem et al. (2018) for COI, H3, 18S, and 28S genes (Table S4). All PCR products were purified with the NucleoSpin^®^ DNA clean-up kit (Faber et al., 2013) and then sent to Macrogen Inc. (Seoul, South Korea) for sequencing for both directions (forward and reverse). The primers used in this study are given in Table S5. Sequences were aligned for each gene region separately using Clustal X v2 (Larkin et al., 2007). Acquired contigs were aligned on BioEdit version 7 (Hall, 2004), edited, and trimmed.

### 2.4. Morphotypes and haplotypes network analyses

The minimum spanning network of morphotypes and of haplotypes were calculated using Arlequin version 3.5.2.2 (Excoffier & Lischer, 2010) and visualized via HapStar (Teacher & Griffiths, 2011).

### 2.5. Bayesian trees

Bayesian trees of COI haplotypes and database mined samples shows the phylogenetic distances: The best model for the MrBayes v3.2 (Ronquist et al., 2012) was chosen via PhyML-SMS v3 software (Lefort et al., 2017). In total four MrBayes runs (two independent for each) were conducted for the haplotypes (H) alone, as well as together with all the database-mined species/samples (DM). The runs were performed based on the general time reversible model with a proportion of invariable sites (GTR+I) for 900’000 (H) and 5’400’000 (DM) combined states with two independent runs. In total, 15 (H) and 45’002 (DM) trees were sampled after discharging a burn-in fraction of 25 % that verified the log likelihood of the cold chain (LnL) stationarity. As the convergence diagnostic, average standard deviation of split frequencies was recorded 0.004 (H) and 0.005 (DM), and Potential Scale Reduction Factors (PSRFs) were close to 1.0 for both (Ronquist et al., 2012). The sister group of *Botrylloides*, the colonial ascidian *Symplegma*, was used as an out-group. The validity of the MCMC chains were confirmed by visual inspection of the LnL distribution to ensure stationarity (Figure S4, C-D). The final trees were visualized with FigTree v.1.4.4

### 2.6. Species analyses

Species delimitation analyses were carried out using the Automatic Barcode Gap Discovery method through the Assemble Species by Automatic Partitioning (ASAP) (Puillandre et al., 2021), a sequence similarity clustering method; and the Poisson Tree Processes (PTP) (Zhang et al., 2013), a tree-based coalescence method. The hypothetical species are defined as Operational Taxonomic Units (OTUs) using these methods. ASAP clusters sequences into partitions consisting of hypothetical species based on the statistical inference of the “barcode gap,” i.e., the gap in the distribution of intra-species and inter-species pairwise distances. ASAP analyses were performed on the web-based interface (Puillandre et al., 2021) (last accessed date: April 2022). Two metric options provided by ASAP for the pairwise distance calculations were used; Jukes-Cantor (JC69) (Jukes & Cantor, 1969) and Kimura 2 parameter (K80) (Kimura, 1980). This strategy allowed excluding possible biases of the selected evolutionary model for the OTU delimitation. PTP analyses were conducted using the Bayesian implementation (bPTP; adds Bayesian support values to delimited species on the input tree), available on the web-based interface (Zhang et al., 2013) (last access date; April 2022). MrBayes trees were generated using 100’000 Markov Chain Monte Carlo (MCMC) generations, subsampling every 100 generations, a burn-in fraction of 0.1 and seeds 123.

### 2.7. Population analyses

Mean COI distances between the NEMS populations and the NCBI database sequences was calculated by a Kimura 2-parameter model (Nei & Kumar, 2000) using MEGA X (Kumar et al., 2018). The Blast suite of NCBI was used to find the matching substitution rates between bases (Johnson et al., 2008).

All population-wide statistics were calculated from the corresponding multiple sequence alignments of the COI locus per population with DnaSP version 6 (Rozas et al., 2017). The following four genetic diversity indices were measured: number of polymorphic sites (N_p_), number of haplotypes (N_h_), nucleotide diversity (π; window length: 100, step size: 25) (Nei & Li, 1979), haplotype diversity (H_d_, window length: 100, step size: 25) (Nei, 1987).

In addition, three associated neutrality test statistics were computed to test the hypothesis that all mutations are selectively neutral: Fu and Li’s D* (F&LD) (Fu & Li, 1993), Fu and Li’s F* (F&LF) (Fu & Li, 1993), Tajima’s D (TajD) (Tajima, 1983).

To compare populations, the pairwise gene flow (N_m_) (Nei, 1973), the pairwise genetic differentiation (F_st_, permutations number: 10’000) (Hudson *et al*., 1992), and the population size changes were calculated with DnaSP. For the population size change, measured population mismatch distributions were compared to expected values for a population with constant population size (Watterson, 1975) using the raggedness statistic, *r* (Harpending, 1994).

All figures were edited with Inkscape (Bah, 2009).

## 3. Results

### 3.1. Morphological records

Overall, 14 different morphotypes were observed among the sampled colonies (Figure 1). Single and double-coloured morphs were recorded: orange-brown, orange, brown-striped, orange-red, white-brown, green-brown, cream-green, red, black-white, pink, cream-violet, orange-striped, red-striped, and brown. These 14 major morphotypes were composed of multiple sub-types with minute differences in the patterns of the colours.

All colonies presented the typical *Botrylloides* ladder-like organization where zooids are aligned side by side while their dorsal lamina was facing the surrounding environment, and the ventral side was located on the attachment side (Figure S1). Four large and four small tentacles were recorded inside the buccal siphon, and eight alternative smaller protrusions were observed (Figure S1). Pigmented blood cells were recorded on the tentacles, especially on the two largest ones. Many pigmented blood cells on both sides of the endostyle were observed over the whole length of the animals.

### 3.2. Life history

The blastogenic cycle of 21 colonies were examined according to the Watanabe (1953) four phase (A-D) staging method. The duration of the cycle varied between 4 to 7 days (Figure S2). In stage A, two primary blastozooids were formed by budding from a single parental zooid. During stage B, blood flow was observed in the cardiac swellings of the primary buds. In stage C, secondary blastozooids were formed from the primary zooids. In stage D, known as take-over, the parental zooids were reabsorbed by the colony while the clonal primary zooids matured (Figure S2).

14 out of the 20 cultured colonies hibernated during the winter season. During hibernation, there was no zooid in the dormant colony. The overall morphology thus consisted of only a carpet-like layer of ampullae (Figure S3). Blood circulation was lower and thicker than usual. Termination of the hibernation was not observed in any colony, even after the end of winter.

The life span and morphology of 20 colonies were documented (Table S6). The average life span of the *B. niger* colonies under the lab conditions was ∼5 months.

### 3.3. Network analysis based on COI

Our samples clustered under four haplotypes; H1, H2, H3, and H4 (Figure 2). The populations from the sampling sites of Kızkalesi, Mezitli, and Konacık were observed to contain more genetic variation, with three unique haplotypes in these regions. No correlation was measured between the genetic and morphological variation for the haplotypes. The highest divergence was in Konacık, where the main haplotype (H1) was separated by 4 mutation steps.

**Figure 2.**
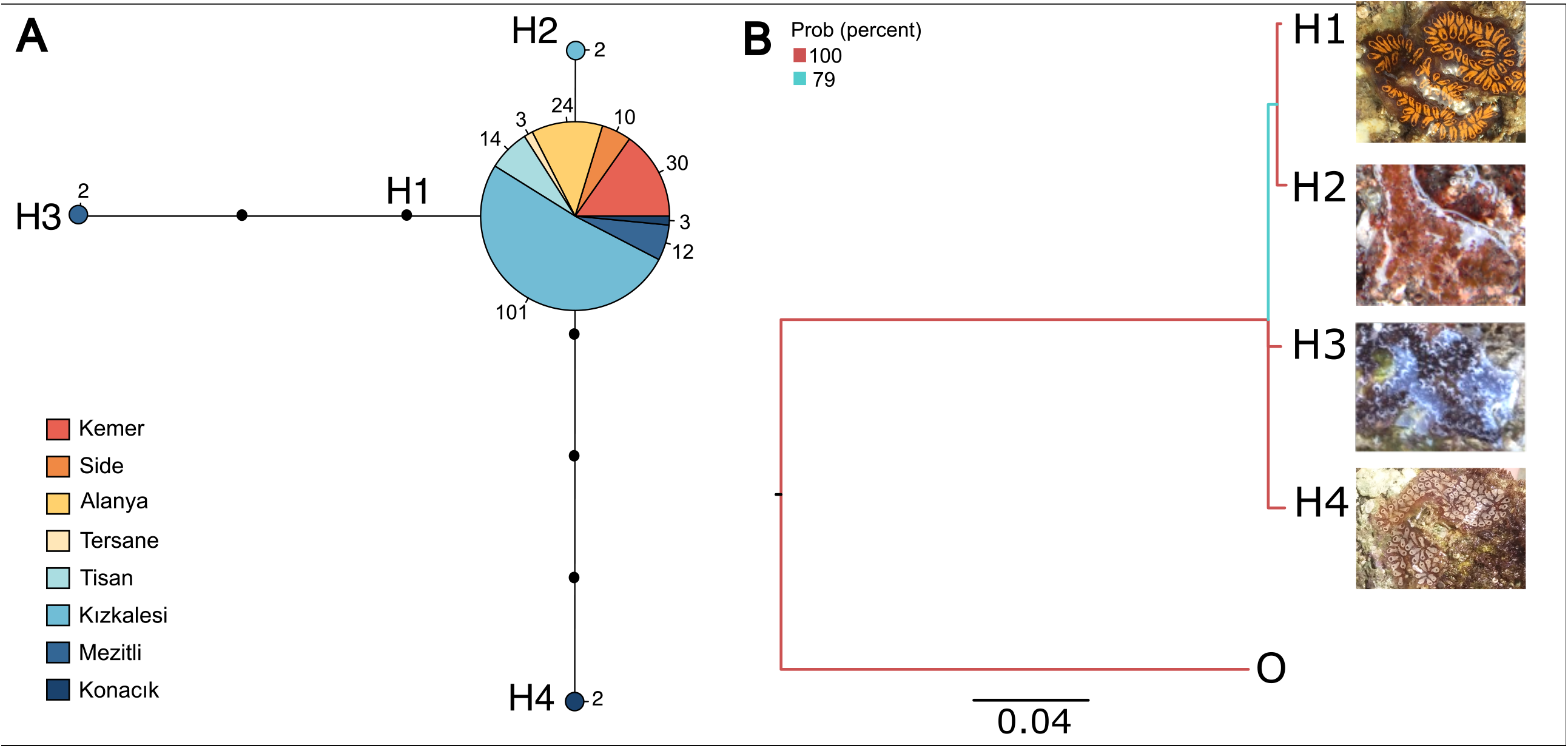
COI haplotypes network of *B. niger* from the NEMS. **A)** Minimum spanning network based on COI haplotypes of *B. niger* locus was constructed. Circles were remarked by the haplotype name. Size differences of circles state frequency while colours reflect regions where the colony was sampled. The region names were given on the left of the network. Numbers represent mutation steps between the haplotypes. **B)** Phylogenetic distance between the COI haplotypes depicted by the best Bayesian tree, together with representative images of each haplotype. The branch colours represent bootstrap value of probability (as percent in the scale). The tree was rooted with the outgroup species *Symplegma brakenhielmi*.

### 3.4. Species delimitation analysis

In total 5 OTUs were assigned for all samples. *Botrylloides cf. lentus* (ON098245_1) was assigned in OTU-1, *Symplegma brakenhielmi* (LS992554_1) in OTU-2, all 27 *Botrylloides niger/nigrum/aff. leachii* reference samples together with all 11 from the present study in OTU-3, two uncharacterized *Botrylloides sp*. samples in OTU-4, and the 54 reference samples of *Botrylloides diegensis/leachii* in OTU-5 (Figure 3). All the main lineages reached 100 % support, with some peripheral linages having lower support down to 60 %. ASAP and PTP results supported the same species clustering (Figure S4).

**Figure 3.**
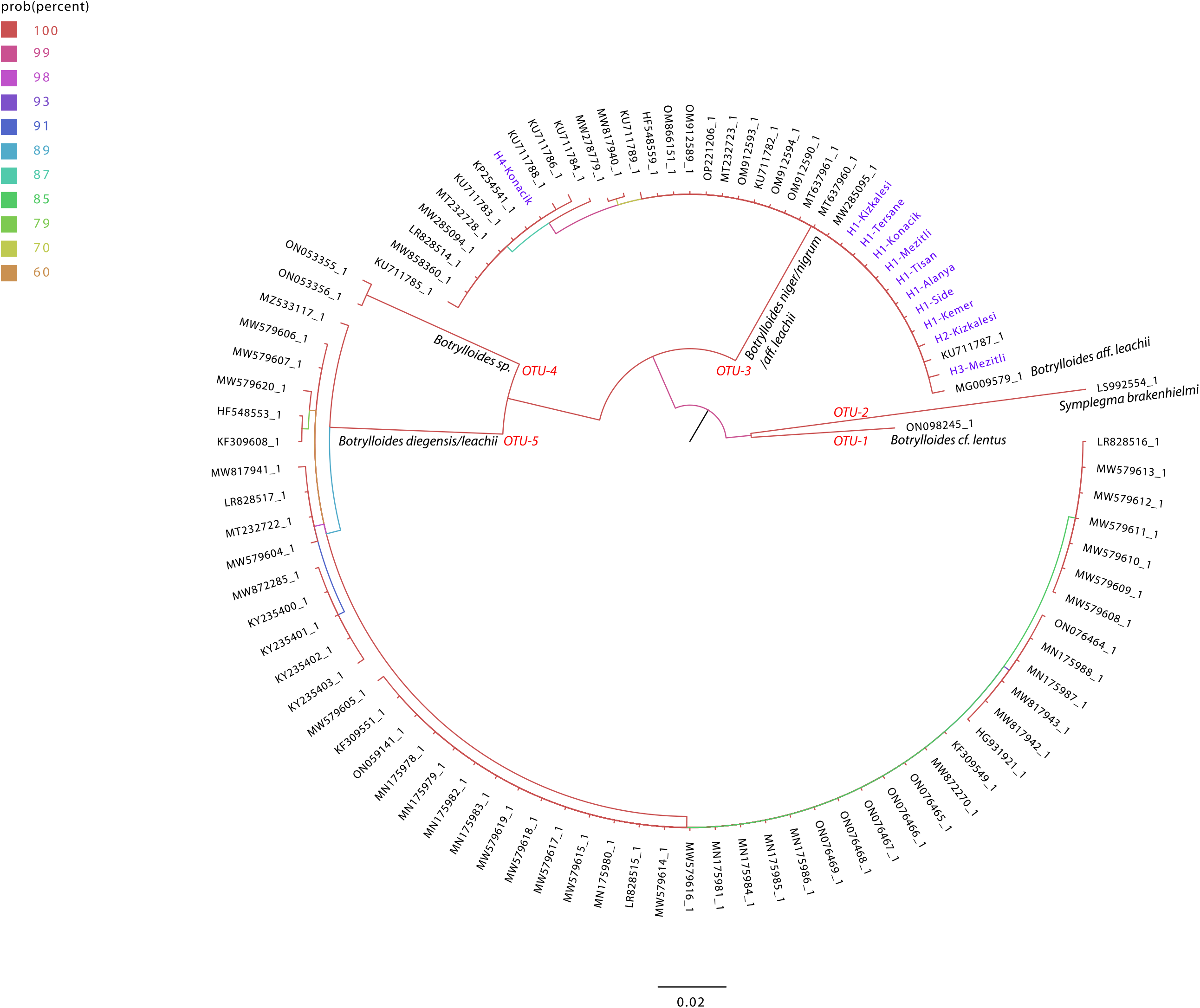
Bayesian majority rule consensus tree reconstructed from the 519 bp COI sequence alignment. Support value is colour-coded as depicted on the left side of the tree. Corresponding OTUs as determined by the ASAP and bPTP analyses are written at the root of each group. The distance scale is given under the tree.

### 3.5. Species assignment by DNA barcoding analysis

The final length of aligned and trimmed COI partial sequences was 519 bp. The divergence among the studied populations was 0.000-0.004 (Table 1). The highest distance was observed between the Konacık and the rest of the NEMS populations. *B. aff. leachii* (MG009579) and *B. niger* (MW858360, LR828514, MW278779) database sequences diverged less than 1% from the present populations. The distances between the NEMS samples and the database reference *B. leachii/B. diegensis* samples were ∼17-23%.

**Table 1.** Pairwise mean COI distances between the NEMS populations and reference populations.

|  | 1 | 2 | 3 | 4 | 5 | 6 | 7 | 8 | 9 | 10 | 11 | 12 | 13 | 14 | 15 | 16 |
| --- | --- | --- | --- | --- | --- | --- | --- | --- | --- | --- | --- | --- | --- | --- | --- | --- |
| 1. Kızkalesi | - |  |  |  |  |  |  |  |  |  |  |  |  |  |  |  |
| 2. Mezitli | 0.001 | - |  |  |  |  |  |  |  |  |  |  |  |  |  |  |
| 3. Tershane | 0.000 | 0.001 | - |  |  |  |  |  |  |  |  |  |  |  |  |  |
| 4. Alanya | 0.000 | 0.001 | 0.000 | - |  |  |  |  |  |  |  |  |  |  |  |  |
| 5. Side | 0.000 | 0.001 | 0.000 | 0.000 | - |  |  |  |  |  |  |  |  |  |  |  |
| 6. Kemer | 0.000 | 0.001 | 0.000 | 0.000 | 0.000 | - |  |  |  |  |  |  |  |  |  |  |
| 7. Tisan | 0.000 | 0.001 | 0.000 | 0.000 | 0.000 | 0.000 | - |  |  |  |  |  |  |  |  |  |
| 8. Konacık | 0.003 | 0.004 | 0.003 | 0.003 | 0.003 | 0.003 | 0.003 | - |  |  |  |  |  |  |  |  |
| 9. BN-IL | 0.000 | 0.001 | 0.000 | 0.000 | 0.000 | 0.000 | 0.000 | 0.003 | - |  |  |  |  |  |  |  |
| 10. BL-IL | 0.004 | 0.005 | 0.004 | 0.004 | 0.004 | 0.004 | 0.004 | 0.007 | 0.004 | - |  |  |  |  |  |  |
| 11. BN-US | 0.005 | 0.005 | 0.005 | 0.005 | 0.005 | 0.005 | 0.005 | 0.005 | 0.005 | 0.008 | - |  |  |  |  |  |
| 12. BN-BR | 0.008 | 0.008 | 0.008 | 0.008 | 0.008 | 0.008 | 0.008 | 0.005 | 0.008 | 0.011 | 0.005 | - |  |  |  |  |
| 13. BL-IT | 0.169 | 0.169 | 0.169 | 0.169 | 0.169 | 0.169 | 0.169 | 0.174 | 0.169 | 0.176 | 0.175 | 0.180 | - |  |  |  |
| 14. BD-FR | 0.196 | 0.195 | 0.196 | 0.196 | 0.196 | 0.196 | 0.196 | 0.201 | 0.196 | 0.192 | 0.202 | 0.209 | 0.008 | - |  |  |
| 15. BL-ES | 0.197 | 0.196 | 0.197 | 0.197 | 0.197 | 0.197 | 0.197 | 0.200 | 0.197 | 0.187 | 0.202 | 0.204 | 0.009 | 0.000 | - |  |
| 16. BL-FR | 0.230 | 0.230 | 0.230 | 0.230 | 0.230 | 0.230 | 0.230 | 0.232 | 0.230 | 0.234 | 0.232 | 0.234 | 0.196 | 0.203 | 0.199 | - |
| 17.Out-group | 0.227 | 0.226 | 0.227 | 0.227 | 0.227 | 0.227 | 0.227 | 0.230 | 0.227 | 0.218 | 0.233 | 0.235 | 0.216 | 0.236 | 0.215 | 0.268 |

Histone 3, 18S rRNA and 28S rRNA sequences from the Kızkalesi station paired at least 99 % with the reference sequences of *B. aff. leachii* (Reem et al., 2018).

### 3.6. Spatial diversity analysis based on COI

The evaluated regional population genetic diversity metrics (Table 2) showed that, despite its lower sample size (n=5), the highest polymorphism (N_p_=4), haplotype diversity (N_h_=0.6), and nucleotide diversity (π=0.0046) were measured in the Konacık population. No polymorphism was documented among the Tisan, Alanya, Side, Tersane, and Kemer populations.

**Table 2.** Genetic diversity indices and neutrality test statistics per NEMS geographical population.

| Populations | n | N<sub>p</sub> | N<sub>h</sub> | π | H<sub>d</sub> | F&LD | F&LF | TajD |
| --- | --- | --- | --- | --- | --- | --- | --- | --- |
| **Kızkalesi** | 38 | 1 | 2 | 0.0001 | 0.053 | -1.758 | -1.823 | -1.129 |
| **Mezitli** | 14 | 3 | 2 | 0.0015 | 0.264 | 1.070 | 0.757 | -0.494 |
| **Tisan** | 14 | 0 | 1 | 0 | 0 | 0 | 0<sup>NS</sup> | 0 |
| **Alanya** | 24 | 0 | 1 | 0 | 0 | 0 | 0<sup>NS</sup> | 0 |
| **Side** | 10 | 0 | 1 | 0 | 0 | 0 | 0<sup>NS</sup> | 0 |
| **Tersane** | 3 | 0 | 1 | 0 | 0 | 0 | 0<sup>NS</sup> | 0 |
| **Kemer** | 30 | 0 | 1 | 0 | 0 | 0 | 0<sup>NS</sup> | 0 |
| **Konacık** | 5 | 4 | 2 | 0.0046 | 0.600 | 1.641 | 1.670 | 1.641 |
| **All** | 138 | 7 | 4 | 0.0004 | 0.071 | 0.257 | -0.557 | -1.857* |

Sequence numbers (n), number of polymorphic sites (N<sub>p</sub>), number of haplotypes (N<sub>h</sub>), nucleotide diversity (π), haplotype diversity (H<sub>d</sub>), Fu and Li's D (F&LD), Fu and Li's F (F&LF) and Tajima's D (TajD). Statistical significance: $p < 0.05$ (\*).

Fu and Li’s D, Fu, and Li’s F, and Tajima’s D tests were calculated to examine the neutrality of mutations (Table 2). While no test was statistically significant at the level of geographical sub-populations, Tajima’s D value for all the NEMS populations resulted in a significant negative value (−1.857, *p* <0.05).

Pairwise gene flow (N_m_) approximations between the eight spatial stations (Table 3) showed the lowest value for the Konacık-Kemer pair (1.58), the highest value for the Kızkalesi-Kemer pair (2422) and no gene flow between the Tisan, Alanya, Side and Kemer samples.

**Table 3.** Pairwise comparison of the gene flow (N_m_) and genetic differentiation (F_st_) between the spatial populations. N_m_ is displayed on the lower triangle and F_st_ on the upper triangle of the comparison table. The statistical significance for F_st_ are indicated.

| $N_m \setminus F_{st}$ | Mezitli | Konacık | Tersane | Tisan | Alanya | Side | Kemer | Kızıkalesi |
| --- | --- | --- | --- | --- | --- | --- | --- | --- |
| **Mezitli** |  | 0.17* | 0.08 | 0.08 | 0.08 | 0.08 | 0.08* | 0.07 |
| **Konacık** | 6.71 |  | 0.25 | 0.25* | 0.25** | 0.25* | 0.25**<br>* | 0.25*** |
| **Tersane** | 8.76 | 3.75 |  | n.c. | n.c. | n.c. | n.c. | 0 |
| **Tisan** | 12 | 1.92 | 0 |  | n.c. | n.c. | n.c. | 0 |
| **Alanya** | 9.16 | 1.66 | 0 | 0 |  | n.c. | n.c. | 0 |
| **Side** | 13.92 | 2.17 | 0 | 0 | 0 |  | n.c. | 0 |
| **Kemer** | 8.29 | 1.58 | 0 | 0 | 0 | 0 |  | 0 |
| **Kızıkalesi** | 16.03 | 2.61 | 7.63 | 96.06 | 576.98 | 46.53 | 2422 |  |

$N_m < 1$ ; there is not enough gene flow to prevent genetic differentiation; $N_m > 1$ , there is enough gene flow to decrease the effects of genetic drift; $N_m > 4$ , populations are involved in randomly mating populations. Statistical significance: $p$ -value $< 0.05$ (\*); $p$ -value $< 0.01$ (\*\*); $p$ -value $< 0.001$ (\*\*\*). n.c.: Not calculated due to a lack of polymorphism.

Pairwise genetic differentiation (F_st_) estimations (Table 3) showed that the Konacık population statistically differed from all the other groups except the Tersane one, and that the Kemer-Mezitli pair also was statistically different.

Population size changes raggedness statistic (*r*) indicated that the only significant result was recorded for the Side population (Figure S5).

### 3.7. Temporal diversity analysis based on COI

The genetic diversity metrics for the Kızkalesi time-series (Table 4) showed two haplotypes and one polymorphic site. The highest nucleotide and haplotype diversities were observed within the November population (π=0.0002, H_d_=0.111), while no diversity was observed for the August-September period. The same polymorphism (N_p_=1) was recorded within the November and the September populations.

**Table 4.** Genetic diversity indices and neutrality test statistics per Kızkalesi temporal population.

| Populations | n | $N_p$ | $N_h$ | $\pi$ | $H_d$ | F&LD | F&LF | TajD |
| --- | --- | --- | --- | --- | --- | --- | --- | --- |
| **11/2017** | 18 | 1 | 2 | 0.0002 | 0.111 | -1.450 | -1.612 | -1.165 |
| **07/2018** | 4 | 0 | 1 | 0 | 0 | 0 | 0 | 0 |
| **08/2018** | 17 | 0 | 1 | 0 | 0 | 0 | 0 | 0 |
| **09/2018** | 26 | 0 | 1 | 0 | 0 | 0 | 0 | 0 |
| **10/2018** | 38 | 1 | 2 | 0.0001 | 0.053 | -1.758 | -1.823 | -1.129 |
| **All** | 103 | 1 | 2 | 0.0001 | 0.038 | 0.491 | 0.080 | -0.912 |

Sequence numbers (n), number of polymorphic sites ( $N_p$ ), number of haplotypes ( $N_h$ ), nucleotide diversity ( $\pi$ ), haplotype diversity ( $H_d$ ), Fu and Li's $D^*$ (F&LD), Fu and Li's $F^*$ (F&LF) and Tajima's $D$ (TajD). No value was measured with a statistically significant $p$ -value $< 0.05$ .

The neutrality test statistics (Table 4) showed no statistically significant values for any population, but negative TajD values for the November and October populations.

Pairwise gene flow (N_m_), pairwise genetic differentiation (F_st_) and population size changes raggedness statistic (*r*) showed no significant value for any of the temporal populations (Figure S6 & Table S7).

## 4. Discussion

In the present study, one primary (COI) and three additional molecular markers (H3, 18S & 28S) were used to identify the genetic diversity of the *B. niger* colonies from the NEMS coasts of Türkiye. Taxonomical assignment based on the morphology of the samples was limited to the genus level (i.e. as a *Botrylloides sp*.) due to the high similarities between the sister species of this taxon (Brunetti, 2009; Reem et al., 2018; Nydam et al., 2021; Salonna et al., 2021). Although the type of larval incubation and the structure of pyloric caecum have been reported to be support species differentiation (Brunetti, 2009), these features are very challenging to assess precisely during punctual field sampling.

Morphological characteristics of the ascidians as a tool for distinct species and genera have remained cryptic, and there is no consensus yet. According to Van Name (1945) and Boyd et al. (1990), morphological variations are accepted to have no taxonomic importance for ascidians. On the other hand, Tarjuelo et al. (2004) suggested that it is possible to differentiate the colour morphs of *Pseudodistoma crucigaster*, a colonial ascidian, based on COI locus. Furthermore, a recent study proposes that although there are morphological overlaps between *Botryllus* and *Botrylloides*, they possess different features (Nydam et al., 2021). While this proposition keeps two genera separated, the morphological separation within *Botrylloides* remains untangled. Aside from the general morphological confusion of *B. niger* with its sister species as *B. leachii, B. diegensis*, or *B. violaceus*, the colour morphs that were given for *Botrylloides simodensis and Botrylloides praelongus* resemble significantly some morphs of the NEMS colonies by highlighting that *Botrylloides* species share highly similar morphotypes (Atsumi & Saito, 2011).

Based our morphological examination, a ladder-like “*leachii* type” zooid organization was found in all NEMS colonies (Brunetti, 2009). At least 14 major morphotypes with various types of striped pigmentation as sub-morphs were recorded within the NEMS. Morphological diversity was similar to the description of Sheets et al. (2016) who indicated 8 different morphotypes for *B. niger* for the 16 worldwide locations. We observed most of the indicated morphotypes in our NEMS colonies, with the addition of a frequent green-violet morph that was not documented in their study. In this study, we assumed that *B. niger* has been introduced to the Mediterranean basin from the Atlantic ocean where it has been proposed to originate from (Sheets et al., 2016). However, the greater number of morphotypes that we have found in the NEMS is challenging this hypothesis, since higher diversity suggests less bottlenecked populations and thus potentially fewer migrations. Similarly, a significant morph variation of *B. schlosseri* were recorded for the Mediterranean colonies highlighting the different adaptive importance based on the pigmentation patterns (Cima et al., 2015). While being a species complex and encompassing a diverse origin from Pacific to Atlantic, Mediterranean colonies of *B. schlosseri* indicated a mixture of native and non-native sub-species (Nydam et al., 2017; Ulman et al., 2017). To understand the origins of the *B. niger*, a broader sampling is thus needed.

The variation in the life history characteristics of colonial ascidians is shaped by their high phenotypic and genetic plasticity to environmental changes (Rinkevich et al., 1993; Blanchoud et al., 2018b). Monitoring the blastogenic cycles of the different morphs showed that for all colonies the blastogenic stages (A-D) were sequential, with a duration that varied from 4 to 7 days. These results are congruent with the previously suggested cycle duration of about one week (Berrill, 1947; Blanchoud et al., 2018a). The shorter cycles are probably related to hibernation (Cima et al., 2015; Hyams et al., 2017). We observed that most of the hibernating colonies in the lab could not recover. Hyams et al. (2017) stated that ∼80% of the hibernating colonies died within five months of the hibernation period. Hibernation seems one of the main reasons for the short life span of the colonies. To understand the hibernation in their natural environment, we recorded the colonial diversity during our samplings in the winter period. We did not observe any colony during the winter in the intertidal zone, where we collected our samples regularly. The measured negative TajD value supports this interpretation of population expansion after a selective sweep, albeit without being statistically significantly. However, Fu and Li’s D* and Fu and Li’s F* non-significant values suggest the opposite process. This apparent contradiction might be due to the absence of colonies during the winter period. When the conditions are not favorable, colonies might migrate to deep waters where the salinity is high enough and settle back to the shores when the conditions are suitable again (Karahan et al., 2016).

Along with the morphological and life history characterization of the NEMS population, we also conducted genetic analyses. Molecular comparisons with the GenBank data showed that the current study sequences match 99-100% with the *B. niger* sequences based on the COI locus; thus, we assigned our species as *Botrylloides niger*. We found five OTUs in based on the species delimitation analyses for the current study and GenBank sequences. OTU-3 consisted of all the *B. niger* haplotypes of the current study together with the GenBank *B. niger/nigrum/a. leachii* sequences. The out-group (*Symplegma brakenhielmi)* and the other *Botrylloides* species, such as *Botrylloides diegensis* were located in separate taxonomic units. We thus support COI as an adequate marker to identify and separate *Botrylloides* species, and *B. niger* in particular. Furthermore, our H3, 18S and 28S sequences matched congruently with the same Israeli *B. aff. leachii* sequences whose COI barcode matched our COI barcode 100% (Reem et al., 2018).

We found four haplotypes from the NEMS based on the COI locus, while Sheets et al. (2016) stated eight COI haplotypes globally for *B. niger*. Considering the coastal zone width, NEMS colonies demonstrate considerable diversity. Besides, the overall haplotype diversity of COI was low (π_COI_: 0.0004), which might stem from a recent bottleneck for the mitochondrial genome or from high introgression rates. Similarly, Sheets et al. (2016) found low diversity for the *B. niger* populations on the Atlantic and Pacific coasts of COI and ANT loci.

Considering the significant negative Tajima’s D values, the null hypothesis indicating that overall the colonies of Turkiye’s NEMS were under negative selection for COI was not rejected. NEMS population thus seems to be under purifying selection, expanding from a restricted population or selective sweep.

Likewise, Sheets et al. (2016) found the *B. niger* populations on the Atlantic coast of Panama, the Pacific coast of Panama, and the coasts of Mexico to be under negative selection for the COI locus, which suggests that a small population size, a founder effect, or a low dispersal. The selective forces acting as the environmental stressors on the mitochondrial DNA might be salinity or temperature (Galtier et al., 2009; Morin et al., 2018; Wei et al., 2019).

The demographic population analyses based on the COI gene have shown that the Side population was under significant subdivision, which could very likely result from a local major freshwater input (the Manavgat stream). Freshwater inputs are known to fluctuate the coastal salinity values. Supportively, Karahan et al. (2016) stated that the dynamics of *B. schlosseri* populations from the California coasts were dramatically affected upon flooding events. Moreover, low-salinity exposed *B. violaceus* larvae were shown to express osmotic stress with an increased mortality rate, suggesting colonial fitness would be reduced due to seasonal storm events that fluctuate seawater salinity (Lambert et al., 2018). Since the lowest salinity record (37.5 ppt) among the sampling stations belongs to Side, the population structure of the *B. niger* colonies seems to be affected by the lower salinity.

COI gene flow estimations were conducted to elucidate the population interactions, but no phylogeographic pattern was recorded. Considering the highest distance between sampling stations (Kemer-Konacık) is approximately 800 kilometers, the larval movement is ecologically restricted. Sea flow and regional maritime traffic possibly affects the larval dispersal from the colonies that are attached to hulls (Lambert, 2001). Nevertheless, the Konacık population seems to be isolated based on its population diversity.

In this study, the population structures and interactions of *B. niger* from the NEMS were investigated using four molecular markers for eight spatial stations and one time-series station. Morphological results showed similar colonial characteristics regarding the blastogenic cycle and hibernation but with a greater number of new morphotype records. Concerning the ambiguities in the classification of *Botrylloides, previously* suggested molecular markers were used to identify the populations from the Turkish coasts of the NEMS. As a result, the COI marker was observed to provide sufficient identification of the species.

## Supplementary materials

Ten supplementary tables and six supplementary figures are provided as supporting information.

## Author contributions

Conceptualization, B.T. and A.K.; Data curation, B.T., E.Ö. and A.K.; Methodology, B.T., E.Ö., S.B. and A.K.; Writing-original draft, B.T., S.B. and A.K. All authors have read and agreed to the published version of the manuscript.

## Institutional review board statement

Not applicable.

## Informed consent statement

Not applicable.

## Data Availability Statement

Sequences, trace files, image files, and the primers information for each COI haplotype were uploaded to the Barcode of Life Data System (Ratnasingham & Hebert, 2007) within the “IMS-METU-Animalia” project file with accession numbers IMS255-19, IMS256-19, and IMS257-19 and BOLD BIN code ACI1328.

The Histone 3, 18S rRNA and 28S rRNA reference sequences of the present study are given in Table S8.

Accession numbers of the COI sequences taken from GenBank are provided in Table S9 (data retrieved from the National Center for Biotechnology Information on 01/09/2022).

Morphotypes of the samples colonies are provided in Table S10.

## Acknowledgments

We thank Dr. Megan Wilson and Dr. Michael Hart for their valuable comments on the final version of the manuscript. We thank Prof. Dr. Baruch Rinkevich, Dr. Jacob Douek, and Guy Paz for their endless support. We thank Oscar Dalkjaer Wigant for proofreading the manuscript. We thank Side and Erdemli Municipality for their kind assistance during our samplings. We also thank the anonymous reviewers for their helpful comments.

