## Supplementary material for "Phylogeographic and morphological analysis of *Botrylloides niger* Herdman, 1886 from the northeastern Mediterranean Sea"

### Supporting Tables

**Table S1.** Details of sampling stations.

| Region | Station | Date | Coordinates | Salinity (ppt) | pH | N |
|---|---|---|---|---|---|---|
| **Antalya** | Kemer | 25/10/2018 | 36°35'-59.55''-N 30°34'-30.66''-E | 40.1 | 7.9 | 30 |
| | Side | 24/10/2018 | 36°46'-00.78''-N 31°23'-07.62''-E | 37.5 | 7.8 | 10 |
| | Alanya | 24/10/2018 | 36°33'-33.85''-N 31°57'-07.41''-E | 39.8 | 8 | 3 |
| | Tersane | 29/10/2018 | 36°32'-05.14''-N 31°59'-53.89''-E | - | - | 3 |
| **Mersin** | Tisan | 03/10/2018 | 36°09'-27.95''-N 33°40'-59.90''-E | 40.2 | 8.35 | 14 |
| | Kızkalesi | 03/10/2018 | 36°27'-27.52''-N 34°08'-38.17''-E | 39-40 | ~ 8 | 37 |
| | Mezitli | 10/4/2018 | 36°43'-58.76''-N 34°31'-18.53''-E | 39.2 | 8.13 | 11 |
| **Hatay** | Konacık | 26/9/2018 | 36°21'-38.79''-N 35°49'-16.90''-E | 40 | 7.9 | 5 |

N: sample number

**Table S2.** Additional spatial samples from the Kızkalesi, Alanya and Mezitli stations.

| Date | N | Station |
|---|---|---|
| 20/4/2012 | 5 | Kızkalesi |
| 27/4/2014 | 26 | Alanya |
| 10/7/2014 | 3 | Kızkalesi |
| 30/9/2014 | 16 | Kızkalesi |
| 11/11/2017 | 3 | Mezitli |
| 10/3/2019 | 1 | Kızkalesi |
| 15/4/2019 | 1 | Kızkalesi |

N: sample number

**Table S3.** Kızkalesi time-series station samplings.

| Date | N |
|---|---|
| November – 2017 | 20 |
| December -2017 | 0 |
| January – 2018 | 0 |
| February – 2018 | 0 |
| March – 2018 | 0 |
| April – 2018 | 0 |
| May – 2018 | 0 |
| June – 2018 | 0 |
| July – 2018 | 6 |
| August – 2018 | 18 |
| September – 2018 | 29 |
| October -2018 | 37 |

N: sample number

**Table S4.** PCR programs for COI, H3, 28S rRNA and 18S rRNA genes.

| PCR program | Denaturation | Annealing | Extension | Final Extension | Cycles number | Reference |
|---|---|---|---|---|---|---|
| DEGCOI | 95ºC (1 min) | 45ºC (30 sec) | 72ºC (1 min) | 72ºC (10 min) | 35 | Reem *et al.*, 2017 |
| H3 | 95ºC (1 min) | 60ºC (45 sec) | 72ºC (1 min) | 72ºC (10 min) | 35 | Reem *et al.*, 2017 |
| 28S | 95ºC (1 min) | 62ºC (45 sec) | 72ºC (1 min) | 72ºC (10 min) | 35 | Reem *et al.*, 2017 |
| 18S | 95ºC (1 min) | 60ºC (45 sec) | 72ºC (1 min) | 72ºC (10 min) | 35 | Reem *et al.*, 2017 |

**Table S5.** Primers used in the present study.

| Primer name | Primer sequence | Product size | Analysed size |
|---|---|---|---|
| DEG COI F2 | 5-'AMWAATCATAAAGATATTRGWAC'-3 | ~700 bp | 519 bp |
| DEG COI R2 | 5-' AARAARGAMGTRTTRAAATTHCGATC'-3 | | |
| H3 F1 | 5-'ATGGCTCGTACCAAGCAGACVGC'-3 | ~350bp | 280 bp |
| H3 R1 | 5-'ATATCCTTRGGCATRATRGTGAC'-3 | | |
| 28S C1 | 5-'ACCCGCTGAATTTAAGCAT'-3 | ~950 bp | 648 bp |
| 28S D2 | 5-' TCCGTGTTTCAAGACGGG'-3 | | |
| 18S A | 5-'AACCTGGTTGATCCTGCCAGT'-3 | ~1750 bp | 966 bp |
| 18S B | 5-'GATCCTTCTGCAGGTTCACCTAC'-3 | | |

**Table S6.** Life history of the 20 colonies cultured in the laboratory. Life cycle corresponds to the mean duration of the blastogenic cycle, measured as the time between two takeovers.

| Colony ID | Sampling site | Sampling date | Morphotype | Haplotype | Culture start day | Life cycle (days) | Culture end date |
|---|---|---|---|---|---|---|---|
| **R1** | Kızkalesi | 18.09.2018 | Orange | H1 | 18.09.2018 | 4 | 12.02.2019 |
| **R4** | Kızkalesi | 18.09.2018 | Brown-striped | H1 | 18.09.2018 | 6 | 16.05.2019 |
| **R5** | Kızkalesi | 18.09.2018 | Cream-violet | H1 | 18.09.2018 | 4 | 16.05.2019 |
| **R6** | Kızkalesi | 18.09.2018 | Green-brown | H1 | 18.09.2018 | 3 | 22.04.2019 |
| **R8** | Kızkalesi | 18.09.2018 | Green-brown | H1 | 18.09.2018 | 5 | 16.05.2019 |
| **R9** | Kızkalesi | 18.09.2018 | Brown-striped | H1 | 18.09.2018 | 5 | 07.12.2018 |
| **GK4** | Kızkalesi | 03.10.2018 | Brown-striped | H1 | 03.10.2018 | 4 | 14.03.2019 |
| **GK7** | Kızkalesi | 03.10.2018 | Brown-striped | H1 | 03.10.2018 | 3 | 01.04.2019 |
| **GK13** | Kızkalesi | 03.10.2018 | Orange-brown | H1 | 03.10.2018 | 4 | 07.12.2018 |
| **GK21** | Kızkalesi | 03.10.2018 | Red | H1 | 03.10.2018 | 5 | 16.05.2019 |
| **G1.R** | Tisan | 03.10.2018 | Cream-violet | H1 | 03.10.2018 | 5 | 25.12.2018 |
| **G1.L** | Tisan | 03.10.2018 | Cream-violet | H1 | 03.10.2018 | 4 | 25.12.2018 |
| **G8** | Tisan | 03.10.2018 | Orange-brown | H1 | 03.10.2018 | 5 | 25.12.2018 |
| **G15** | Tisan | 03.10.2018 | Green-brown | H1 | 03.10.2018 | 5 | 26.02.2019 |
| **L15** | Konacık | 26.09.2018 | White-brown | H4 | 26.09.2018 | 5 | 16.05.2019 |
| **L18** | Konacık | 26.09.2018 | Brown-striped | H1 | 26.09.2018 | 4 | 16.05.2019 |
| **Z1** | Kızkalesi | 03.08.2018 | Cream-violet | H1 | 03.08.2018 | 6 | 14.03.2019 |
| **Z2** | Kızkalesi | 03.08.2018 | Orange | H1 | 03.08.2018 | 5 | 15.02.2019 |
| **Z4** | Kızkalesi | 03.08.2018 | Cream-violet | H1 | 03.08.2018 | 6 | 14.03.2019 |
| **NN** | Kızkalesi | 03.08.2018 | Red | H1 | 03.08.2018 | 5 | 26.02.2019 |

**Table S7.** Pairwise comparison of the gene flow ($N_m$) and genetic differentiation ($F_{st}$) between the temporal populations of Kızkalesi. $N_m$ is displayed on the lower triangle and $F_{st}$ on the upper triangle of the comparison table.

| $N_m$ \ $F_{st}$ | 11/2017 | 07/2018 | 08/2018 | 09/2018 | 11/2018 |
|---|---|---|---|---|---|
| **11/2017** | - | 0 | 0 | 0 | -0.037 |
| **07/2018** | 16.16 | - | n.c. | n.c. | 0 |
| **08/2018** | 21420 | 0 | - | n.c. | 0 |
| **09/2018** | 643 | 0 | 0 | - | 0 |
| **10/2018** | -32.79 | 11.04 | 161.13 | 878.22 | - |

$N_m$ <1: there is not enough gene flow to prevent genetic differentiation; $N_m$ > 1: there is enough gene flow to decrease the effects of genetic drift; $N_m$ > 4: populations are involved in randomly mating populations. No statistical significance was measured for $F_{st}$. n.c.: Not calculated due to a lack of polymorphism .

**Table S8.** *B. niger* H3, 18S and 28S reference sequences used in the present study.

| Gene | Sequence |
|---|---|
| **H3** | GCTCCACGAAAGCAGCTCGCCACAAAGGCGGCCAGGAAGAGCGCGCCAGCCACGGGCGGCGTCA<br>AGAAGCCTCATCGTTACAGGCCGGGCACGGTGGCGCTCCGTGAGATCAGACGGTACCAAAAGTCC<br>ACCGAGCTGCTCATACGCAAGCTTCCGTTCCAGCGACTGGTGCGCGAGATCGCCCAGGACTTCAA<br>GACCGACCTTCGCTTCCAGAGCAGCTCCGTGATGGCGTTGCAAGAAGCCAGCGAAGCCTACCTCG<br>TGGGCCTGTTCGAAGACACCA |
| **18S** | CTGCGAATGGCTCATTAAATCAGTCTTGGTTTATTTGGTCTCGAGAGCGAAGGTGGATAACTGTGG<br>CAATTCCAGAGCTAATACATGCAATTAGCGCCGACTTCGGGAGGCGTGCTTTTATTGGATCAAAAC<br>CGACCGGGTTCGCCCGTCTCTTTTGATGACTCTGGATAACCACGCGGATCGCGCGGTCTTGTGCCG<br>GCGACAAACCATTCAAGTGTCTGACCTATCAACTTTCGAAGGTAAGCTACGGGCTTACCTTTGTGA<br>TAACGGGTGACGGGGAATCAGGGTTCGATTCCGGAGAGGGAGCCTGAGAAACGGCTACCACATC<br>CAAGGAAGGCAGCAGGCGCGCAAATTACCCATTCCCGACACGGGGAGGTAGTGACGAAAAATAA<br>CAATACAGGACTCTAACGAGGCCCTGTAATTGGAATGAGTACATTCTAAAACTCTTAACGAGTAT<br>CCATTGGAGGGCAAGTCTGGTGCCAGCAGCCGCGGTAATTCCAGCTCCAAAAGTGTATGCTAAAG<br>TTGTTGCGGTTGAAAAGCTCGTAGTTGGATATTGGGCGAGCGCGGTCGGTCCGTCGCAGGGCGTG<br>TACTGGTCGCGTTCGTCTCGAGCTTCGGTTCTCCGTCGGTGCTCTTGACTGAGTGTCGGCGGTGGC<br>CGAGAAGTTTACTTTGAAAAAATTAGAGTGTTCAAAGCAGGCTGGTCGCCTGAATAGTGTTGCAT<br>GGAATAATGGAATAGGACCTCGGTTCTATTTTGTTGGTTTTCGGAACGGAGGTAATGATTAAGAG<br>GGACAGACGGGGGTGTCCGTACTCTGCCGTTAGAGGTGAAATTCTTGGATCGGCGGAAGACGAAC<br>TACTGCGAAAGCATTCACCAAGAATGTTTTCTTTAATCAAGAGCGAAAGTCAGAGGTTCGAAGAC<br>GATCAGATACCGTCCTAGTTCTGACTATAAACGATGCCAACTAGCGATCGG |
| **28S** | CAACGGGGATTCCCCGAGTAACGGCGAGTGAATCGGGAACAGTCCAGCGCTGAATCTGCACGTTC<br>TTGAAGCGTGCCGAGTTGTGGCGTAAGGAAGTCTCTGTGCGGTCGTCGGCGAGCGTCGCTGTCCTT<br>CTGATCGAGGCCTCATCCCGGTGCGGGTGTCAGGCCCATAAGGGCGCTTGCCGCGGCCGCTTCGA<br>GTCTTCCCGGAGTCGGGTTGTTTGAGAATGCAGCCTAAAGCGGGTGGTAAACTCCACCTAAGACT<br>AAATACGGTCGCGAGACCGATAGCGAACAAGTACCGTGAGGGAAAGTTGAAAAGCACTTTGGAG<br>AGAGAGTTCAAAAGTACGTGAAACCGTCAAGAGGCAAACGGGAGAGCCCGTCCGGGCTCGGGTA<br>CGCTTTCAGTTGGGCGCGGAAGCGCGCTGCAGGCAGTTGTGCGGACGCTCGCGCGCCGTCTCTGTC<br>GTAGCACGCACGCCGCTCCAGCGCACTAGCGTCCCGAGCAGGCCACGACCGGTTCGAAGTCGGCC<br>AGAAGCGCCGCGCGAAGGTGACCCACGCTTCGGCGGGGGTGTTACAGCGCGCGTGCGTCGAGGCC<br>CGGCTACGGATCGAGGTTACGAGTCCGTGCGCGCCTTCCTTCGGGAGGGTTGGCTGCGAGCGT |

**Table S9.** Accession numbers of the GenBank COI reference sequences.

| Scientific Name | Accession number | Table 1 group |
|---|---|---|
| *Botryllus schlosseri* | AY600987.1 | Out-group |
| *Botrylloides leachii* | HF548553.1 | |
| *Botrylloides nigrum* | HF548559.1 | BN-IL |
| *Botrylloides leachii* | HG931921.1 | |
| *Botrylloides leachii* | KF309549.1 | |
| *Botrylloides leachii* | KF309551.1 | |
| *Botrylloides leachii* | KF309608.1 | |
| *Botrylloides leachii* | KF309644.1 | BL-ES |
| *Botrylloides nigrum* | KP254541.1 | |
| *Botrylloides nigrum* | KU711782.1 | |
| *Botrylloides nigrum* | KU711783.1 | |
| *Botrylloides nigrum* | KU711784.1 | |
| *Botrylloides nigrum* | KU711785.1 | |
| *Botrylloides nigrum* | KU711786.1 | |
| *Botrylloides nigrum* | KU711787.1 | |

| Scientific Name | Accession number | Table 1 group |
|---|---|---|
| *Botrylloides nigrum* | KU711788.1 | |
| *Botrylloides nigrum* | KU711789.1 | |
| *Botrylloides leachii* | KY235400.1 | |
| *Botrylloides leachii* | KY235401.1 | |
| *Botrylloides leachii* | KY235402.1 | |
| *Botrylloides leachii* | KY235403.1 | |
| *Botrylloides niger* | LR828514.1 | BN-BR |
| *Botrylloides leachii* | LR828515.1 | |
| *Botrylloides leachii* | LR828516.1 | |
| *Botrylloides leachii* | LR828517.1 | |
| *Symplegma brakenhielmi* | LS992554.1 | |
| *Botrylloides leachii* | MG009578.1 | BL-IT |
| *Botrylloides aff. leachii* | MG009579.1 | BL-IL |
| *Botrylloides leachii* | MK978812.1 | BL-FR |
| *Botrylloides diegensis* | MN076483.1 | BD-FR |
| *Botrylloides diegensis* | MN175978.1 | |
| *Botrylloides diegensis* | MN175979.1 | |
| *Botrylloides diegensis* | MN175980.1 | |
| *Botrylloides diegensis* | MN175981.1 | |
| *Botrylloides diegensis* | MN175982.1 | |
| *Botrylloides diegensis* | MN175983.1 | |
| *Botrylloides diegensis* | MN175984.1 | |
| *Botrylloides diegensis* | MN175985.1 | |
| *Botrylloides diegensis* | MN175986.1 | |
| *Botrylloides diegensis* | MN175987.1 | |
| *Botrylloides diegensis* | MN175988.1 | |
| *Botrylloides diegensis* | MT232722.1 | |
| *Botrylloides niger* | MT232723.1 | |
| *Botrylloides niger* | MT232728.1 | |
| *Botrylloides niger* | MT637960.1 | |
| *Botrylloides niger* | MT637961.1 | |
| *Botrylloides nigrum* | MW278779.1 | BN-US |
| *Botrylloides niger* | MW285094.1 | |
| *Botrylloides niger* | MW285095.1 | |
| *Botrylloides diegensis* | MW579604.1 | |
| *Botrylloides diegensis* | MW579605.1 | |
| *Botrylloides diegensis* | MW579606.1 | |
| *Botrylloides diegensis* | MW579607.1 | |
| *Botrylloides diegensis* | MW579608.1 | |
| *Botrylloides diegensis* | MW579609.1 | |
| *Botrylloides diegensis* | MW579610.1 | |
| *Botrylloides diegensis* | MW579611.1 | |

| Scientific Name | Accession number | Table 1 group |
|---|---|---|
| *Botrylloides diegensis* | MW579612.1 | |
| *Botrylloides diegensis* | MW579613.1 | |
| *Botrylloides diegensis* | MW579614.1 | |
| *Botrylloides diegensis* | MW579615.1 | |
| *Botrylloides diegensis* | MW579616.1 | |
| *Botrylloides diegensis* | MW579617.1 | |
| *Botrylloides diegensis* | MW579618.1 | |
| *Botrylloides diegensis* | MW579619.1 | |
| *Botrylloides diegensis* | MW579620.1 | |
| *Botrylloides niger* | MW817940.1 | |
| *Botrylloides diegensis* | MW817941.1 | |
| *Botrylloides diegensis* | MW817942.1 | |
| *Botrylloides diegensis* | MW817943.1 | |
| *Botrylloides niger* | MW858360.1 | BN-US |
| *Botrylloides diegensis* | MW872270.1 | |
| *Botrylloides diegensis* | MW872285.1 | |
| *Botrylloides diegensis* | MZ533117.1 | |
| *Botrylloides niger* | OM866151.1 | |
| *Botrylloides niger* | OM912589.1 | |
| *Botrylloides niger* | OM912590.1 | |
| *Botrylloides niger* | OM912593.1 | |
| *Botrylloides niger* | OM912594.1 | |
| *Botrylloides sp.* | ON053355.1 | |
| *Botrylloides sp.* | ON053356.1 | |
| *Botrylloides diegensis* | ON059141.1 | |
| *Botrylloides diegensis* | ON076464.1 | |
| *Botrylloides diegensis* | ON076465.1 | |
| *Botrylloides diegensis* | ON076466.1 | |
| *Botrylloides diegensis* | ON076467.1 | |
| *Botrylloides diegensis* | ON076468.1 | |
| *Botrylloides diegensis* | ON076469.1 | |
| *Botrylloides cf. lentus* | ON098245.1 | |
| *Botrylloides niger* | OP221206.1 | |

**Table S10**. Correspondance between sample IDs and the determined haplotypes.

| Sample ID | Haplotype |
|---|---|
| BN1-NOV17 | H2 |
| BN2-NOV17 | H1 |
| BN3-NOV17 | H1 |
| BN4-NOV17 | H1 |
| BN5-NOV17 | H1 |
| BN6-NOV17 | H1 |
| BS1-NOV17 | H1 |
| BS2-NOV17 | H1 |
| BS3-NOV17 | H1 |
| BS4-NOV17 | H1 |
| BS5-NOV17 | H1 |
| BS6-NOV17 | H1 |
| BV1-NOV17 | H1 |
| BV2-NOV17 | H1 |
| BV3-NOV17 | H1 |
| BV4-NOV17 | H1 |
| BV5-NOV17 | H1 |
| BV8-NOV17 | H1 |
| GK1-OCT18 | H1 |
| GK2-OCT18 | H1 |
| GK3-OCT18 | H1 |
| GK4-OCT18 | H1 |
| GK5-OCT18 | H1 |
| GK6-OCT18 | H1 |
| GK7-OCT18 | H1 |
| GK8-OCT18 | H1 |
| GK9-OCT18 | H1 |
| GK10-OCT18 | H1 |
| GK11-OCT18 | H1 |
| GK12-OCT18 | H1 |
| GK13-OCT18 | H1 |
| GK14-OCT18 | H1 |
| GK15-OCT18 | H1 |
| GK17-OCT18 | H1 |
| GK18-OCT18 | H1 |
| GK20-OCT18 | H1 |
| GK21-OCT18 | H1 |
| GK22-OCT18 | H1 |

| Sample ID | Haplotype |
|---|---|
| GK24-OCT18 | H1 |
| GK25-OCT18 | H1 |
| GK26-OCT18 | H2 |
| GK27-OCT18 | H1 |
| GK29-OCT18 | H1 |
| GK30-OCT18 | H1 |
| GK31-OCT18 | H1 |
| GK32-OCT18 | H1 |
| GK33-OCT18 | H1 |
| GK35-OCT18 | H1 |
| GK38-OCT18 | H1 |
| GK39-OCT18 | H1 |
| GK40-OCT18 | H1 |
| GK41-OCT18 | H1 |
| GK43-OCT18 | H1 |
| GK44-OCT18 | H1 |
| GK45-OCT18 | H1 |
| GK46-OCT18 | H1 |
| J18-BS2-JULY18 | H1 |
| J18-BS7-JULY18 | H1 |
| J18-BS8-JULY18 | H1 |
| J18-BS9-JULY18 | H1 |
| R1-SEP18 | H1 |
| R2-SEP18 | H1 |
| R4-SEP18 | H1 |
| R5-SEP18 | H1 |
| R6-SEP18 | H1 |
| R7-SEP18 | H1 |
| R8-SEP18 | H1 |
| R9-SEP18 | H1 |
| R10-SEP18 | H1 |
| R11-SEP18 | H1 |
| R12-SEP18 | H1 |
| R14-SEP18 | H1 |
| R15-SEP18 | H1 |
| R16-SEP18 | H1 |
| R21-SEP18 | H1 |
| R28-SEP18 | H1 |
| R29-SEP18 | H1 |

| Sample ID | Haplotype |
|---|---|
| R32-SEP18 | H1 |
| R36-SEP18 | H1 |
| R37-SEP18 | H1 |
| R39-SEP18 | H1 |
| R40-SEP18 | H1 |
| R41-SEP18 | H1 |
| R42-SEP18 | H1 |
| R43-SEP18 | H1 |
| R44-SEP18 | H1 |
| Z1-AUG18 | H1 |
| Z2-AUG18 | H1 |
| Z3-AUG18 | H1 |
| Z4-AUG18 | H1 |
| Z5-AUG18 | H1 |
| Z6-AUG18 | H1 |
| Z8-AUG18 | H1 |
| Z9-AUG18 | H1 |
| Z10-AUG18 | H1 |
| Z11-AUG18 | H1 |
| Z12-AUG18 | H1 |
| Z14-AUG18 | H1 |
| Z15-AUG18 | H1 |
| Z16-AUG18 | H1 |
| Z19-AUG18 | H1 |
| Z20-AUG18 | H1 |
| Z21-AUG18 | H1 |
| M2-12-Mezitli | H3 |
| M2-17-Mezitli | H3 |
| L9-Konacik | H4 |
| L15-Konacik | H4 |
| K1-Tersane | H1 |
| K2-Tersane | H1 |
| K3-Tersane | H1 |
| L2-Konacik | H1 |
| L17-Konacik | H1 |
| L18-Konacik | H1 |
| M2-6-Mezitli | H1 |
| M2-7-Mezitli | H1 |
| M2-10-Mezitli | H1 |

| **Sample ID** | **Haplotype** |
|---|---|
| M2-13-Mezitli | H1 |
| M2-14-Mezitli | H1 |
| M2-15-Mezitli | H1 |
| M2-19-Mezitli | H1 |
| M2-20-Mezitli | H1 |
| M2-22-Mezitli | H1 |
| Mz_1-Mezitli | H1 |
| Mz_2-Mezitli | H1 |
| Mz_3-Mezitli | H1 |
| G1-Tisan | H1 |
| G2-Tisan | H1 |
| G4-Tisan | H1 |
| G5-Tisan | H1 |
| G6-Tisan | H1 |
| G8-Tisan | H1 |
| G9-Tisan | H1 |
| G10-Tisan | H1 |
| G11-Tisan | H1 |
| G12-Tisan | H1 |
| G13-Tisan | H1 |
| G14-Tisan | H1 |
| G15-Tisan | H1 |
| G16-Tisan | H1 |
| A1-Alanya | H1 |
| A1_i-Alanya | H1 |
| A3_i-Alanya | H1 |
| A4_i-Alanya | H1 |
| A6-Alanya | H1 |
| A7_i-Alanya | H1 |
| A8-Alanya | H1 |
| A8_i-Alanya | H1 |
| A9_i-Alanya | H1 |
| A10_i-Alanya | H1 |
| A10_i_2-Alanya | H1 |
| A11-Alanya | H1 |
| A12-Alanya | H1 |
| A14-Alanya | H1 |
| A15-Alanya | H1 |
| A16-Alanya | H1 |

| Sample ID | Haplotype |
|---|---|
| A20-Alanya | H1 |
| A23-Alanya | H1 |
| A24-Alanya | H1 |
| A25-Alanya | H1 |
| A26-Alanya | H1 |
| C10-Alanya | H1 |
| C16-Alanya | H1 |
| C17-Alanya | H1 |
| C18-Side | H1 |
| C19-Side | H1 |
| C20-Side | H1 |
| C22-Side | H1 |
| C24-Side | H1 |
| C25-Side | H1 |
| C26-Side | H1 |
| C27-Side | H1 |
| C28-Side | H1 |
| C29-Side | H1 |
| C30-Kemer | H1 |
| C31-Kemer | H1 |
| C32-Kemer | H1 |
| C33-Kemer | H1 |
| C34-Kemer | H1 |
| C35-Kemer | H1 |
| C38-Kemer | H1 |
| C39-Kemer | H1 |
| C40-Kemer | H1 |
| C41-Kemer | H1 |
| C42-Kemer | H1 |
| C43-Kemer | H1 |
| C44-Kemer | H1 |
| C45-Kemer | H1 |
| C46-Kemer | H1 |
| C47-Kemer | H1 |
| C48-Kemer | H1 |
| C49-Kemer | H1 |
| C50-Kemer | H1 |
| C51-Kemer | H1 |
| C52-Kemer | H1 |

| Sample ID | Haplotype |
|---|---|
| C54-Kemer | H1 |
| C55-Kemer | H1 |
| C56-Kemer | H1 |
| C58-Kemer | H1 |
| C59-Kemer | H1 |
| C60-Kemer | H1 |
| C61-Kemer | H1 |
| C62-Kemer | H1 |
| C63-Kemer | H1 |
| GK1-Kizkalesi | H1 |
| GK2-Kizkalesi | H1 |
| GK3-Kizkalesi | H1 |
| GK4-Kizkalesi | H1 |
| GK5-Kizkalesi | H1 |
| GK6-Kizkalesi | H1 |
| GK7-Kizkalesi | H1 |
| GK8-Kizkalesi | H1 |
| GK9-Kizkalesi | H1 |
| GK10-Kizkalesi | H1 |
| GK11-Kizkalesi | H1 |
| GK12-Kizkalesi | H1 |
| GK13-Kizkalesi | H1 |
| GK14-Kizkalesi | H1 |
| GK15-Kizkalesi | H1 |
| GK17-Kizkalesi | H1 |
| GK18-Kizkalesi | H1 |
| GK20-Kizkalesi | H1 |
| GK21-Kizkalesi | H1 |
| GK22-Kizkalesi | H1 |
| GK24-Kizkalesi | H1 |
| GK25-Kizkalesi | H1 |
| GK26-Kizkalesi | H1 |
| GK27-Kizkalesi | H1 |
| GK29-Kizkalesi | H1 |
| GK30-Kizkalesi | H1 |
| GK31-Kizkalesi | H1 |
| GK32-Kizkalesi | H1 |
| GK33-Kizkalesi | H1 |
| GK35-Kizkalesi | H1 |

| Sample ID | Haplotype |
|---|---|
| GK38-Kizkalesi | H1 |
| GK39-Kizkalesi | H1 |
| GK40-Kizkalesi | H1 |
| GK41-Kizkalesi | H1 |
| GK43-Kizkalesi | H1 |
| GK44-Kizkalesi | H1 |
| GK45-Kizkalesi | H1 |
| GK46-Kizkalesi | H1 |

**Supporting Figures**

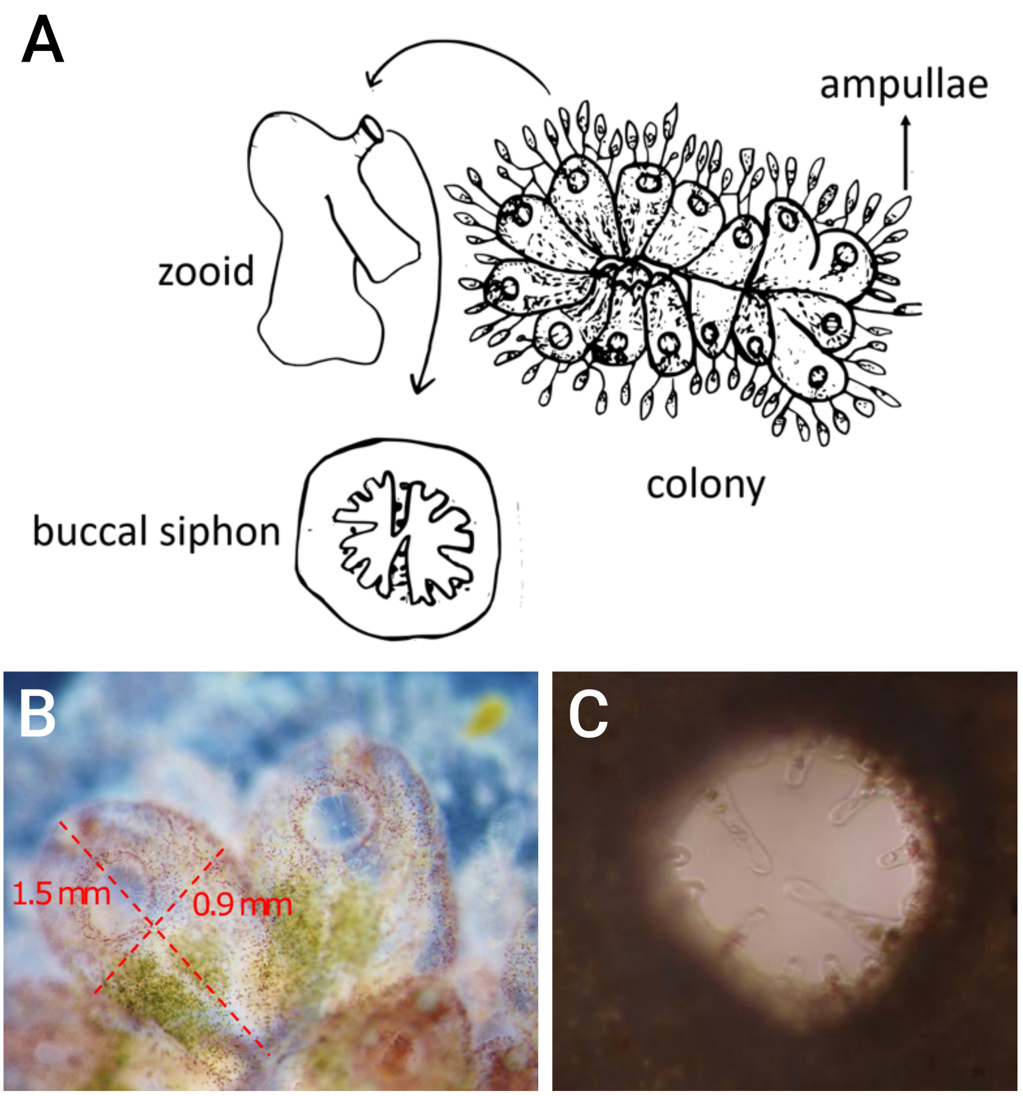


**Figure S1.** Morphology of *Botrylloides niger*. **A**) An illustration of *B. niger* colonial system, highlighting key anatomical features including colony, zooid, ampullae (vascular termini), and buccal siphon. **B**) Top view of two zooid overlaid with measurements of its size. **C**) Magnified picture of a buccal siphon showing its tentacles.

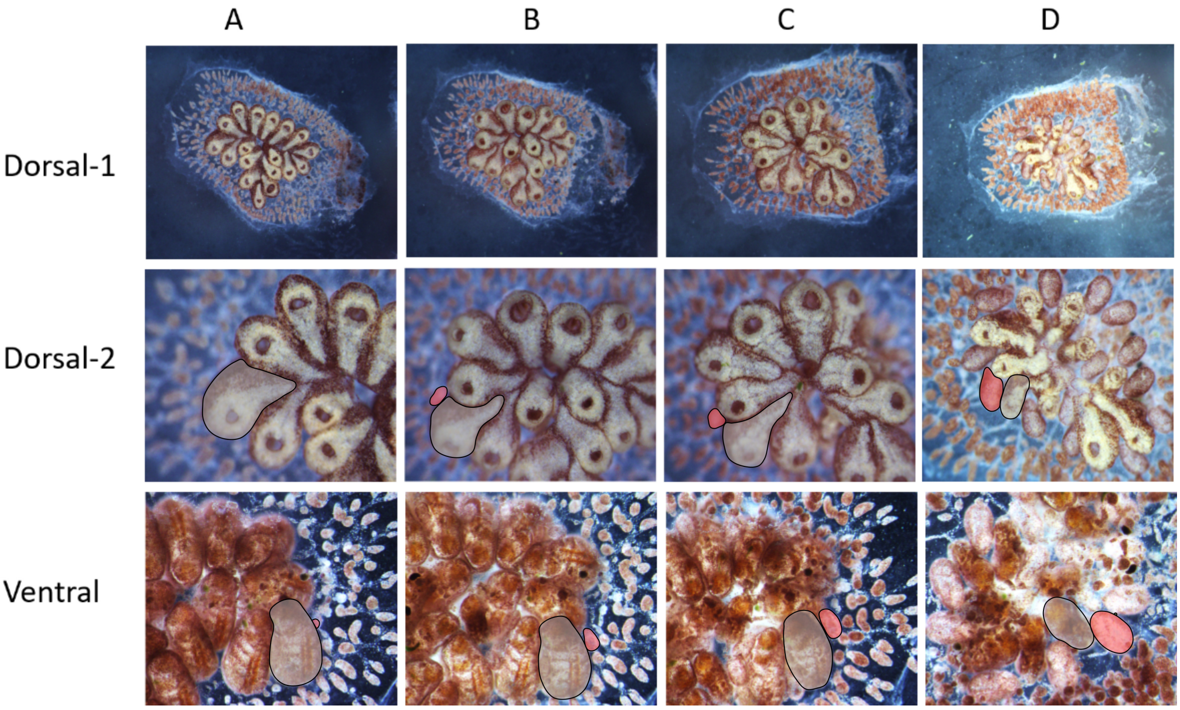


**Figure S2.** The blastogenic cycle of *B. niger*. The cycle was monitored on a daily basis, both from the dorsal and the ventral side of the colony. The letters above each column indicate the corresponding blastogenic stage. New parental zooids are growing new primary buds in the stage A. These new buds are increasing in size and developing secondary buds at stage B and further in stage C. The parental zooids are absorbed at stage D and the primary zooids are replacing them as the new parental zooids in the stage A of the upcoming cycle. The outline of a zooid (grey), of one of its primary bud (pink) and of one of its secondary bud (green) are depicted.

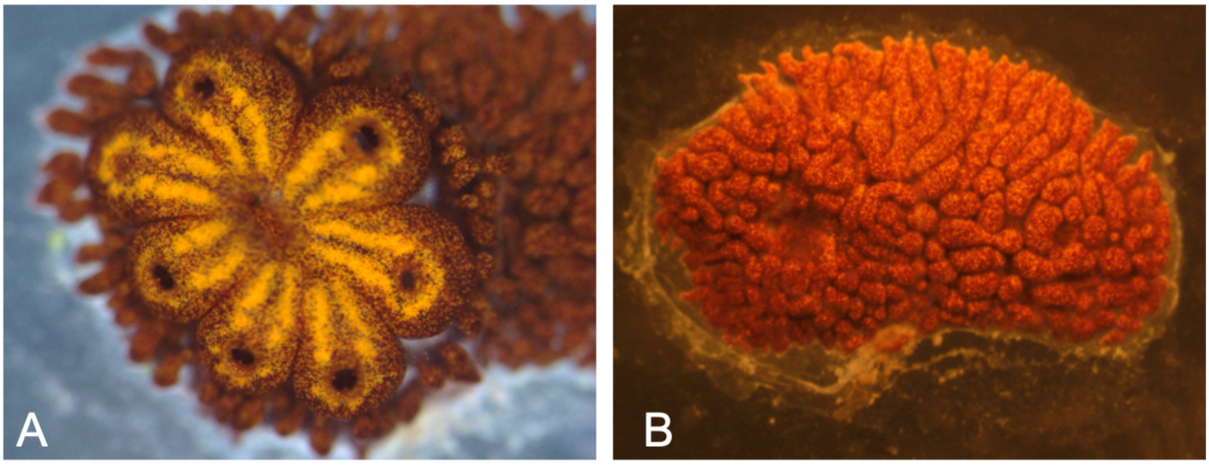


**Figure S3.** Hibernation of *B. niger*. **A**) A common *B. niger* morph during the active filter feeding phase. Seven zooids are visible with their siphons open. **B**) The same colony during hibernation. No zooid is present, only a dense mat of highly pigmented ampullae are visible inside the tunic.

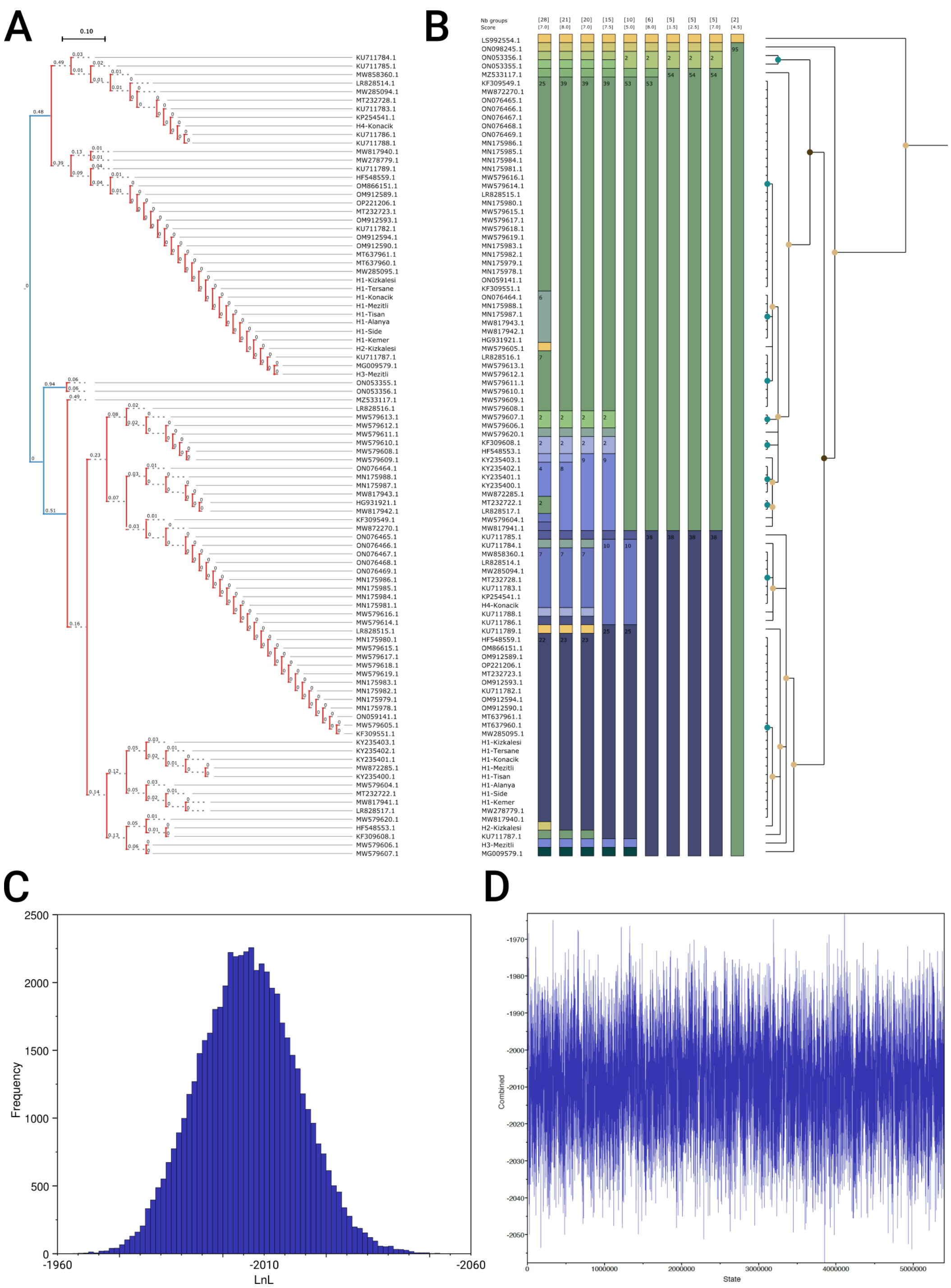


**Figure S4.** Delimiting Operational Taxonomic Units (OTUs) and Bayesian tracer. **A)** The tree-based coalescence method from the Poisson Tree Processes (PTP). Blue lines represent different OTUs, red line the same OTUs. **B)** Assemble Species by Automatic Partitioning (ASAP) score: The colors are representing different OTUs. Number inside each OTU represent the total number of samples assigned to that OTU (one is not indicated). Ten partitioning of increasing granularity (from right to left) are depicted, with their associated ASAP score (the lower the better). **C)** An example of a stationary MCMC chain displaying a Gaussian distribution of its LnL posterior probabilities, after the discarded 0.1 burn-in fraction. **D)** Traces of LnL values for two independent stationary runs combining 5400000 iterations.

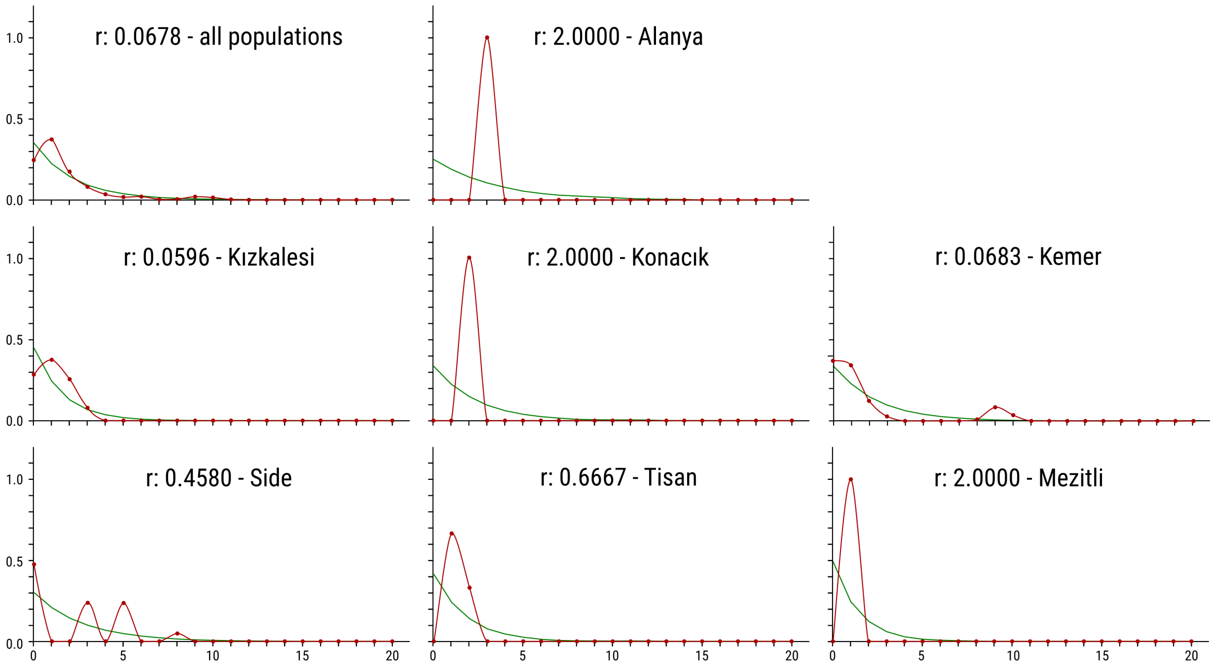


**Figure S5.** Population size changes analysis based on the mismatch distribution of the COI gene for Kızkalesi, Mezitli, Konacık, and all populations. Raggedness statistic values (*r*) for each analysis was supplied ($r > 0.05$). The X-axis is the number of pairwise differences while the Y-axis represents frequency. The red line indicates the observed frequency, and the green line the expected frequency for a stable population.

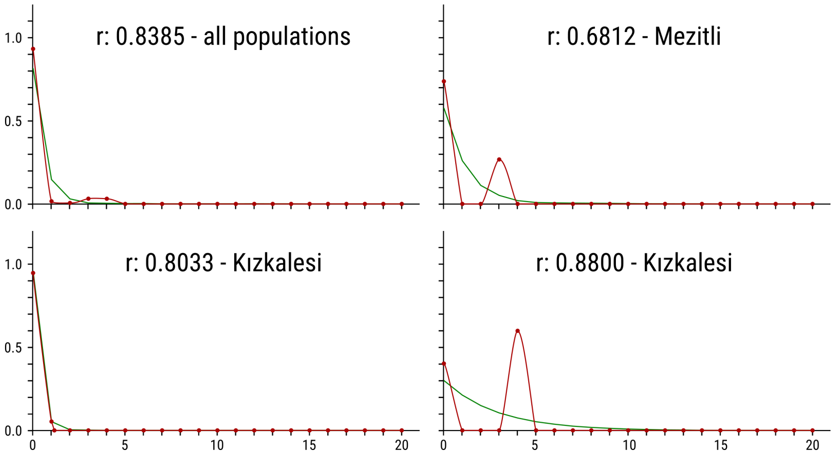


**Figure S6.** Population size changes of Kızkalesi time-series station based on mismatch distribution of COI. Raggedness statistic values (*r*) for each analysis was supplied ($r > 0.05$). The X-axis is the number of pairwise differences while the Y-axis represents frequency. The red line indicates the observed frequency, and the green line the expected frequency for a stable population.
